# Functional and structural characterization of AtAbf43C: An exo-1,5-⍺-L-arabinofuranosidase from *Acetivibrio thermocellus* DSM1313

**DOI:** 10.1101/2025.04.06.647456

**Authors:** Joey L. Galindo, Philip D. Jeffrey, Angela Zhu, A. James Link, Jonathan M. Conway

## Abstract

The *Acetivibrio thermocellus* DSM1313 genome codes for seven predicted glycoside hydrolase family 43 (GH43) enzymes, four of which remain uncharacterized. This study describes the function and structure of one such enzyme, AtAbf43C, from GH43 subfamily 26 (GH43_26) which acts as an ⍺-L-arabinofuranosidase (EC 3.2.1.55). AtAbf43C is active on para-nitrophenol-⍺-L-arabinofuranoside (pNPAra), with optimal activity observed at pH 5.5 and 65 ℃. Multiple crystal structures of AtAbf43C were obtained, in which an N-terminal carbohydrate binding module family 42 (CBM42) domain displays a ß-trefoil type fold and the C-terminal GH43 domain displays a canonical 5-bladed ß-propeller motif. One structure, which was solved with two L-arabinofuranose molecules bound to ß- and ɣ-subdomains of the CBM42, builds upon previous literature suggesting the ⍺-binding pocket of the AtAbf43C CBM42 is non-functional. Furthermore, structural alignment with the substrate bound structure of a closely homologous GH43_26 exo-⍺-1,5-arabinofuranosidase, SaAraf43A from *Streptomyces avermitilis* (PDB 3AKH), allowed for identification of the conserved catalytic triad via site-directed mutagenesis in AtAbf43C, as well as insight into the deep-narrow topology of the AtAbf43C binding pocket that suggested it would be active on similar arabino-oligosaccharide (AOS) substrates as SaAraf43A. Subsequent liquid chromatography-mass spectrometry (LC-MS) analysis of polysaccharides and oligosaccharides hydrolyzed by AtAbf43C provides experimental evidence confirming this enzyme acts in an exo manner primarily towards ⍺-1,5 linked arabino-oligosaccharides.

## Introduction

Plant cell walls are comprised of large biopolymers, primarily cellulose, hemicellulose and lignin, which form the basis for lignocellulosic biomass (1–4). Efficient degradation of the various components of lignocellulose has important applications in many bioprocesses, most notably for the sustainable bioproduction of fuels and chemicals from renewable lignocellulosic feedstocks, but also in commercial food and beverage processing (2, 3, 5). Complete deconstruction of lignocellulose requires a diverse array of Carbohydrate Active enZymes (CAZymes) (3–8). The domains from these CAZymes have been categorized into various families by homology in the CAZy database (7). Glycoside hydrolase (GH) domains, which cleave *O*-glycosidic bonds, are the dominant catalytic players in lignocellulose degradation (4, 6–9). Additionally, Carbohydrate Binding Module (CBM) domains are a type of non-catalytic domain, commonly encoded in the same protein as catalytic CAZy domains, that bind to carbohydrate substrates and heavily influence enzymatic activity and specificity (4, 6–9).

⍺-L-arabinofuranosidases (EC 3.2.1.55), a type of GH enzyme which hydrolyze terminal non-reducing ⍺-1,2, ⍺-1,3, or ⍺-1,5 linked arabinofuranose residues, play an important role in the degradation of arabinose-containing hemicelluloses such as arabinoxylan, arabinan, and arabinogalactan (3–5, 10). Owing to the diversity of these polysaccharides, ⍺-L-arabinofuranosidases vary widely in their specificities towards a given substrate. Some ⍺-L-arabinofuranosidases are only active on small substrates such as short arabino-oligosaccharides (AOS) or arabino-xylo-oligosaccharides (AXOS) as well as the synthetic compound para-nitrophenol-α-L-arabinofuranoside (pNPAra), while other ⍺-L-arabinofuranosidases are active primarily on large polysaccharides like arabinan or arabinoxylan (3, 5, 10, 11). Additionally, a subset of this latter category of ⍺-L-arabinofuranosidases, also known as arabinoxylan arabinofuranohydrolases, specifically cleave arabinofuranosyl residues from arabinoxylans (3, 5, 10, 11). ⍺-L-arabinofuranosidases can be found across several GH families including GH2, 3, 43, 51, 54, 62, and 159 (3, 5). The large GH family 43, which was recently divided into 37 subfamilies, commonly contains galactan 1,3-β-galactosidases (EC 3.2.1.145), β-xylosidases (EC 3.2.1.37), and endo-⍺-L arabinanases (EC 3.2.1.99), in addition to ⍺-L-arabinofuranosidases (8, 12, 13). Several families of CBMs are also commonly found associated with GH43 ⍺-L-arabinofuranosidases, most notably CBM6 and 42 (8, 12–14). CBM6 domains commonly bind directly to polysaccharides like xylan and amorphous cellulose, while CBM42 domains typically recognize small arabino-oligosaccharides or the terminal non-reducing arabinofuranosyl residues of arabinan or arabinoxylan (8, 13, 14).

*Acetivibrio thermocellus* (basionym: *Clostridium thermocellum*) is a thermophilic Gram-positive obligate anaerobic bacterium that has been heavily studied for its ability to efficiently break down lignocellulose, by natively producing a variety of cellulose and hemicellulose degrading enzymes (2, 4, 12, 15). To date three ⍺-L-arabinofuranosidases from *A. thermocellus* have been characterized including an intracellular GH51 family ⍺-L-arabinofuranosidase active towards AOS and AXOS, and two extracellular cellulosomal GH43 ⍺-L-arabinofuranosidases (4, 12, 16, 17). The first of these GH43 ⍺-L-arabinofuranosidases (*Ct*43Araf; subfamily 16) was most active to arabinoxylan polysaccharides, while the second (AxB8; subfamily 29) acted as a bi-functional β-xylosidase/⍺-L-arabinofuranosidase with primary activity towards small AXOS (4, 12, 17). Furthermore, of the seven predicted GH43 genes in *A. thermocellus* genome, only three have been characterized including the two aforementioned ⍺-L-arabinofuranosidases as well as a 1,3-β-galactosidase (1,3Gal43A; subfamily 24) (4, 12, 17, 18). Additionally, three of the four uncharacterized *A. thermocellus* GH43 proteins contain CBM42 domains (4, 14). Previously Ribeiro *et al*. expressed these three CBM42 domains as isolated truncations, and tested their binding of various natural substrates, finding they most strongly bind arabinoxylan and arabinan (14).

In this work we describe the unique crystal structure and function of a fourth *A. thermocellus* GH43 ⍺-L-arabinofuranosidase (AtAbf43C; subfamily 26) that acts primarily in an exo manner towards the ⍺-1,5 linkages present in arabinan and smaller AOS substrates rather than those of arabinoxylan and associated AXOS. We solved the crystal structure of the full enzyme and each of its domains including a structure of its CBM42 bound to arabinose. We characterized the optimal activity of AtAbf43C on a variety of substrates and demonstrated that its activity is exo in nature with AOSs as its primary substrate. Taken together our work provides new insight into the structure and function of a thermophilic ⍺-L-arabinofuranosidase.

## Results

### Sequence Analysis and Diversity of Characterized GH43_26 Enzymes

The gene encoding the AtAbf43C protein in *A. thermocellus* DSM1313 (Locus Tag: *Clo1313_2794*; GenBank Protein accession #: ADU75776.1) has an open reading frame of 1743 bp and is identical in sequence to the *Cthe_2138* gene in *A. thermocellus* ATCC27405. The predicted 580 amino acid protein consists of a N-terminal signal peptide (residues 1-19), a CBM42 domain (residues 29-160), a GH43 domain (residues 179-488), and a C-terminal dockerin I domain (residues 511-568) (**Fig. 1a**). This suggested *Clo1313_2794* likely codes for a secreted enzyme that is part of the extracellular *A. thermocellus* cellulosome. The top non-identical BLASTp hits (67-93% identity) to AtAbf43C are comprised of almost all GH43 enzymes from other *Acetivibrio* species, many of which are predicted to be putative β-xylosidases or ⍺-L-arabinofuranosidases (Table S1). Ten bacterial GH43_26 enzymes (Table S2) cataloged in the CAZy database have previously been characterized (19–25). All of these GH43_26 enzymes are ⍺-1,5-arabinofuranosidases active towards AOS or arabinan. Individually these enzymes have 49-63% identity to AtAbf43C and a phylogenetic tree constructed after a multiple protein sequence alignment shows their diversity (**Fig. 1b**; Table S2).

**Figure 1.**
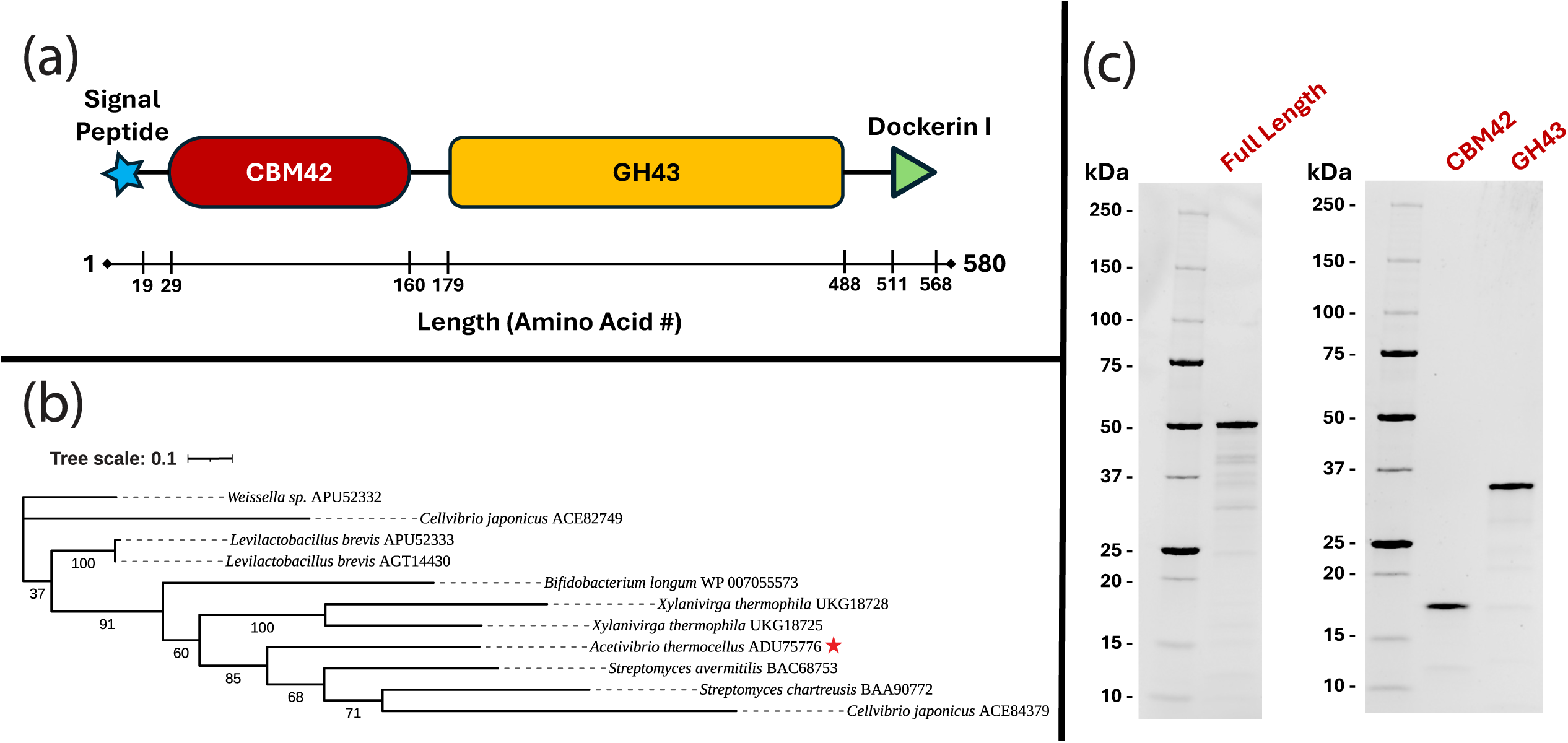
(a) The general architecture of the *Clo1313_2794* gene. **(b)** A phylogenetic tree constructed after a multiple protein sequence alignment using ClustalOmega of the GenBank protein ascension # (ADU75776.1) corresponding to *Clo1313_2794* (★) with the other characterized members of the GH43_26 subfamily. **(c**) Protein gel of the purified full length AtAbf43C protein and its truncated versions: CtAbd43C_CBM42 and AtAbf43C_GH43.

### AtAbf43C Enzymatic Activity

To determine the biochemical properties of AtAbf43C, recombinant AtAbf43C (Residues 21-511, lacking the signal peptide and dockerin I domain) and truncation mutants AtAbf43C_CBM42 (Residues 21-167) and AtAbf43C_GH43 (Residues 167-494) were expressed and purified (**Fig. 1a & c; Table S3**). Using *para*-nitrophenol-⍺-L-arabinofuranoside (pNPAra) as the substrate, we investigated the optimal reaction conditions for AtAbf43C. AtAbf43C showed optimal activity at pH 5.5 and 65 ℃, though the enzyme retained >75% of its maximum activity within range of pH 5-6.5 and a temperature of 55-70 ℃ (**Fig. 2a & b**). When incubated at elevated temperatures, AtAbf43C retains at least 46-48% of its initial activity and at least 28% after 6 hours when incubated at 55-60 ℃ and 65 ℃ respectively (**Fig. 2c**). This is consistent with the results of the temperature optimization assay in which enzyme activity begins to dimmish at 70 ℃ and above (**Fig. 2b**). The effects of additives on the activity of AtAbf43C were also tested (**Fig. 2d**). Some divalent ions, including Ca^2+^, Mg^2+^, and Co^2+^, appeared to have a positive effect on enzymatic activity, with Ca^2+^ addition having the largest effect. In contrast, Zn^2+^ and Cu^2+^ caused a significant decrease in activity. The effects observed with other additives were not statistically significant (p-value ≤ 0.05) (**Table S4**).

**Figure 2.**
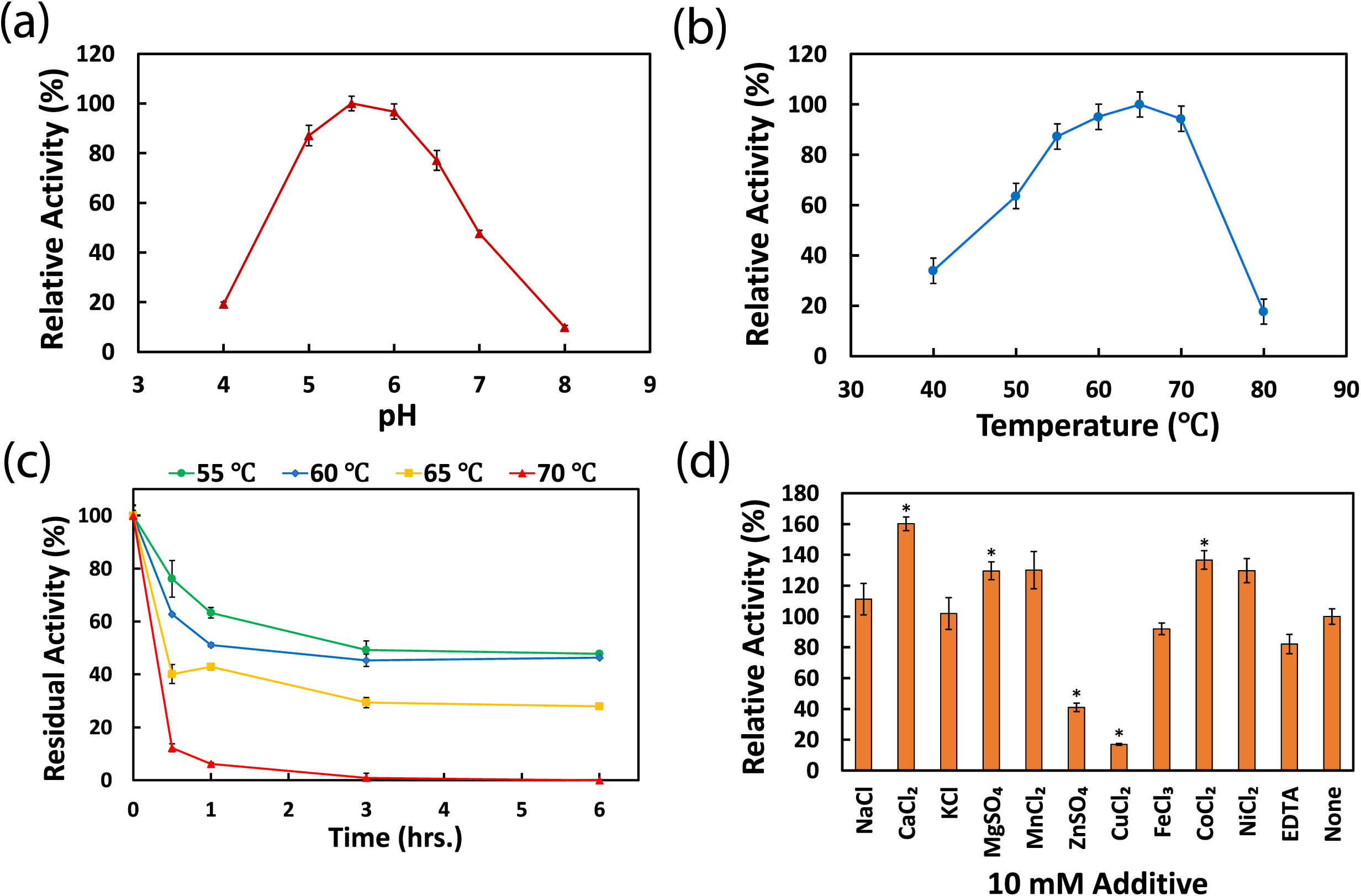
Effect of environmental conditions on AtAbf43C activity on pNPAra. Error bars represent standard deviations between triplicate technical replicates at each reaction condition. (**a)** pH optimization performed at 55°C. **(b)** Temperature optimization performed at pH 5.5. **(c)** Thermostability test of *CtAbf34C.* **(d)** Effect of various 10 mM additives on enzymatic activity. Asterisks indicate statistical significance (P≤0.05) compared with the untreated condition.

### Substrate Specificity

At the optimal pH and temperature AtAbf43C and AtAbf43C_GH43 had specific activities of 4.95±0.28 U/mg and 5.67±0.63 U/mg respectively on pNPAra, while AtAbf43C_CBM42 showed no activity on pNPAra, with U defined as mM/s of released *para*-nitrophenol (pNP) (**Table S5a**). This confirmed that AtAbf43C_GH43 contained the active catalytic domain. In addition to pNPAra, the activity of AtAbf43C and its truncated versions were tested on several other substrates. These included natural substrates: wheat arabinoxylan (WAX), beechwood xylan (BX), and sugar beet arabinan (SBA), which were tested using the dinitrosalicylic acid (DNS) reducing assay; as well as three additional synthetic pNP glycosides: para-nitrophenol-ß-D-xylopyranoside (pNPXy), para-nitrophenol-⍺-D-galactopyranoside (pNP⍺Gal), and para-nitrophenol-ß-D-galactopyranoside (pNPßGal). However, no activity on any of these other substrates was detected (**Table S5a & b**). Kinetic parameters were then determined at optimal conditions on pNPAra for AtAbf43C and AtAbf43C_GH43 (**Table 1, Table S6**). Full substrate saturation could not be achieved for either protein, as the *K_m_* values were so high as to approach the solubility limit of pNPAra in aqueous solution.

**Table 1.** Kinetic parameters determined for AtAbf43C and AtAbf43C_GH43 on pNPAra at optimal <u>conditions.</u>

| Protein | $K_m$ (mM) | $k_{cat}$ ($s^{-1}$) | $k_{cat}/K_m$ ($s^{-1} mM^{-1}$) |
| --- | --- | --- | --- |
| AtAbf43C | 22.7±4.1 | 269.2±25.5 | 11.9±2.4 |
| AtAbf43C_GH43 | 59.6±12.1 | 237.5±33.4 | 4.0±1.0 |

### Crystal Structure and Mutagenesis Study of AtAbf43C

Crystal structures were obtained for the full-length AtAbf43C protein (PDB code: 9NXG) at a resolution of 1.32Å as well as individual domains AtAbf43C_CBM42 (PDB code: 9NXI) and AtAbf43C_GH43 (PDB code: 9NXJ) at 1.75 Å and 2.32Å respectively. Additionally, a 1.75Å crystal structure (PDB code: 9NXH) was solved for AtAbf43C soaked in L-arabinose immediately prior to freezing and mounting in which two L-arabinofuranose molecules were found bound to the CBM42 domain. A summary of refinement statistics for all 4 structures can be found in **Table 2**.

**Table 2.**
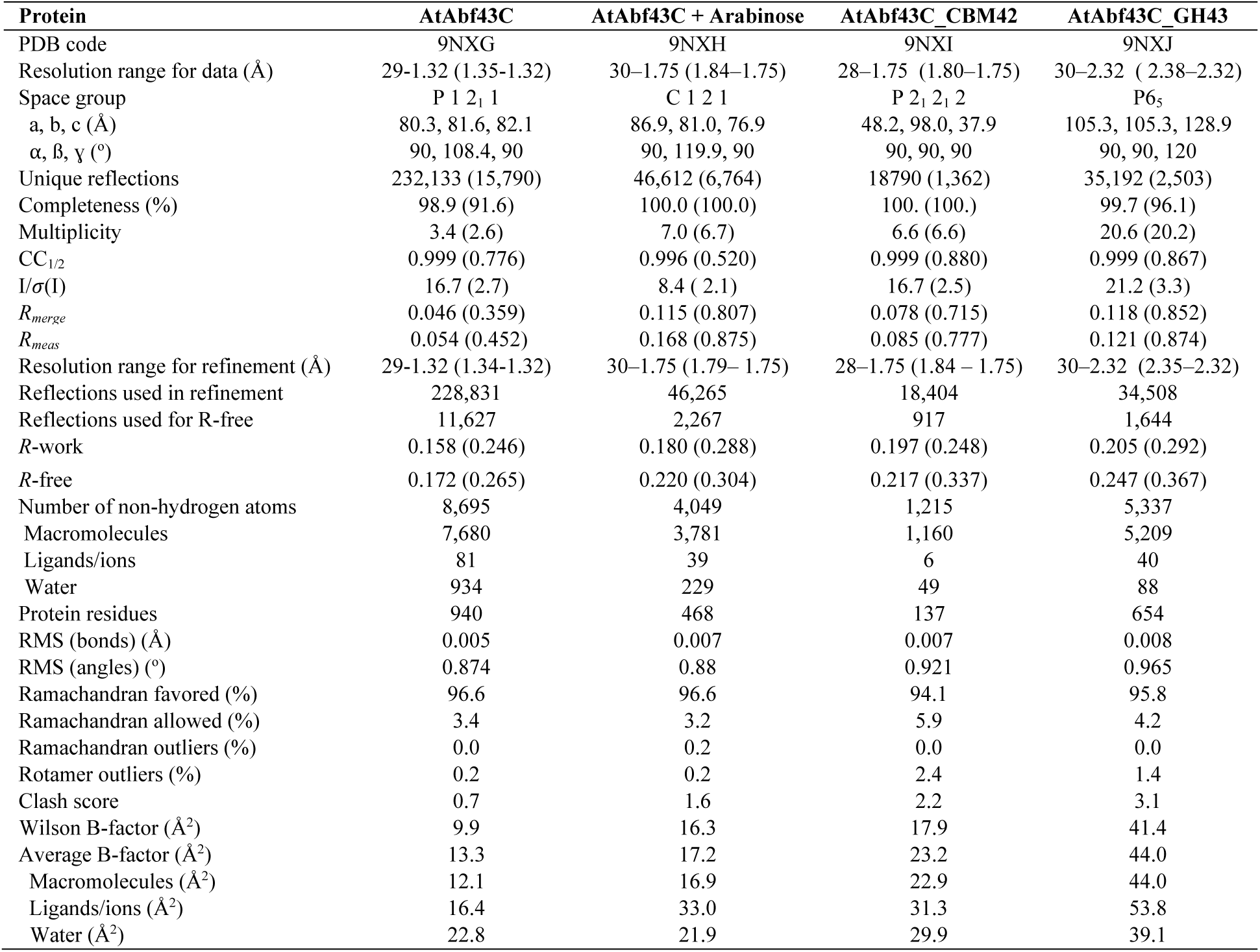
Summary of data and refinement statistics for the structures of AtAbf43C.

| Protein | AtAbf43C | AtAbf43C + Arabinose | AtAbf43C_CBM42 | AtAbf43C_GH43 |
| --- | --- | --- | --- | --- |
| PDB code | 9NXG | 9NXH | 9NXI | 9NXJ |
| Resolution range for data (Å) | 29-1.32 (1.35-1.32) | 30-1.75 (1.84-1.75) | 28-1.75 (1.80-1.75) | 30-2.32 (2.38-2.32) |
| Space group | P 1 2<sub>1</sub> 1 | C 1 2 1 | P 2<sub>1</sub> 2<sub>1</sub> 2 | P6<sub>5</sub> |
| a, b, c (Å) | 80.3, 81.6, 82.1 | 86.9, 81.0, 76.9 | 48.2, 98.0, 37.9 | 105.3, 105.3, 128.9 |
| α, β, γ (°) | 90, 108.4, 90 | 90, 119.9, 90 | 90, 90, 90 | 90, 90, 120 |
| Unique reflections | 232,133 (15,790) | 46,612 (6,764) | 18790 (1,362) | 35,192 (2,503) |
| Completeness (%) | 98.9 (91.6) | 100.0 (100.0) | 100. (100.) | 99.7 (96.1) |
| Multiplicity | 3.4 (2.6) | 7.0 (6.7) | 6.6 (6.6) | 20.6 (20.2) |
| CC<sub>1/2</sub> | 0.999 (0.776) | 0.996 (0.520) | 0.999 (0.880) | 0.999 (0.867) |
| I/σ(I) | 16.7 (2.7) | 8.4 (2.1) | 16.7 (2.5) | 21.2 (3.3) |
| R<sub>merge</sub> | 0.046 (0.359) | 0.115 (0.807) | 0.078 (0.715) | 0.118 (0.852) |
| R<sub>meas</sub> | 0.054 (0.452) | 0.168 (0.875) | 0.085 (0.777) | 0.121 (0.874) |
| Resolution range for refinement (Å) | 29-1.32 (1.34-1.32) | 30-1.75 (1.79-1.75) | 28-1.75 (1.84-1.75) | 30-2.32 (2.35-2.32) |
| Reflections used in refinement | 228,831 | 46,265 | 18,404 | 34,508 |
| Reflections used for R-free | 11,627 | 2,267 | 917 | 1,644 |
| R-work | 0.158 (0.246) | 0.180 (0.288) | 0.197 (0.248) | 0.205 (0.292) |
| R-free | 0.172 (0.265) | 0.220 (0.304) | 0.217 (0.337) | 0.247 (0.367) |
| Number of non-hydrogen atoms | 8,695 | 4,049 | 1,215 | 5,337 |
| Macromolecules | 7,680 | 3,781 | 1,160 | 5,209 |
| Ligands/ions | 81 | 39 | 6 | 40 |
| Water | 934 | 229 | 49 | 88 |
| Protein residues | 940 | 468 | 137 | 654 |
| RMS (bonds) (Å) | 0.005 | 0.007 | 0.007 | 0.008 |
| RMS (angles) (°) | 0.874 | 0.88 | 0.921 | 0.965 |
| Ramachandran favored (%) | 96.6 | 96.6 | 94.1 | 95.8 |
| Ramachandran allowed (%) | 3.4 | 3.2 | 5.9 | 4.2 |
| Ramachandran outliers (%) | 0.0 | 0.2 | 0.0 | 0.0 |
| Rotamer outliers (%) | 0.2 | 0.2 | 2.4 | 1.4 |
| Clash score | 0.7 | 1.6 | 2.2 | 3.1 |
| Wilson B-factor (Å<sup>2</sup>) | 9.9 | 16.3 | 17.9 | 41.4 |
| Average B-factor (Å<sup>2</sup>) | 13.3 | 17.2 | 23.2 | 44.0 |
| Macromolecules (Å<sup>2</sup>) | 12.1 | 16.9 | 22.9 | 44.0 |
| Ligands/ions (Å<sup>2</sup>) | 16.4 | 33.0 | 31.3 | 53.8 |
| Water (Å<sup>2</sup>) | 22.8 | 21.9 | 29.9 | 39.1 |

The full length AtAbf43C arabinose-soaked structure (PDB 9NXH) consists of the smaller N-terminal CBM42 domain connected via a short 19 amino acid linker to the larger C-terminal catalytic GH43 domain (**Fig. 3a**). In addition to the two arabinose molecules bound to the CBM42 domain, the structure also contains three glycerol molecules, and a central magnesium ion in the GH43 domain. The catalytic GH43 domain displays the 5-bladed ß-propeller fold typical of GH43 enzymes (11, 13, 24–30) (**Fig. 3b**). The GH43 domain of AtAbf43C does not have a C-terminal ß-jelly roll domain found in some subfamilies of GH43 proteins, however, the N-terminal strand of the domain appears to form part of the 5^th^ blade in the blade V structure in what is colloquially termed a “molecular velcro” (11, 25, 26, 28, 30) (**Fig. 3b**). This closure in the structure, thought to provide extra structural stability, is not found in many GH43 enzymes in which the 5^th^ blade in the beta propeller only consists of residues found in the C-terminal strand (11, 27, 28, 30).

**Figure 3.**
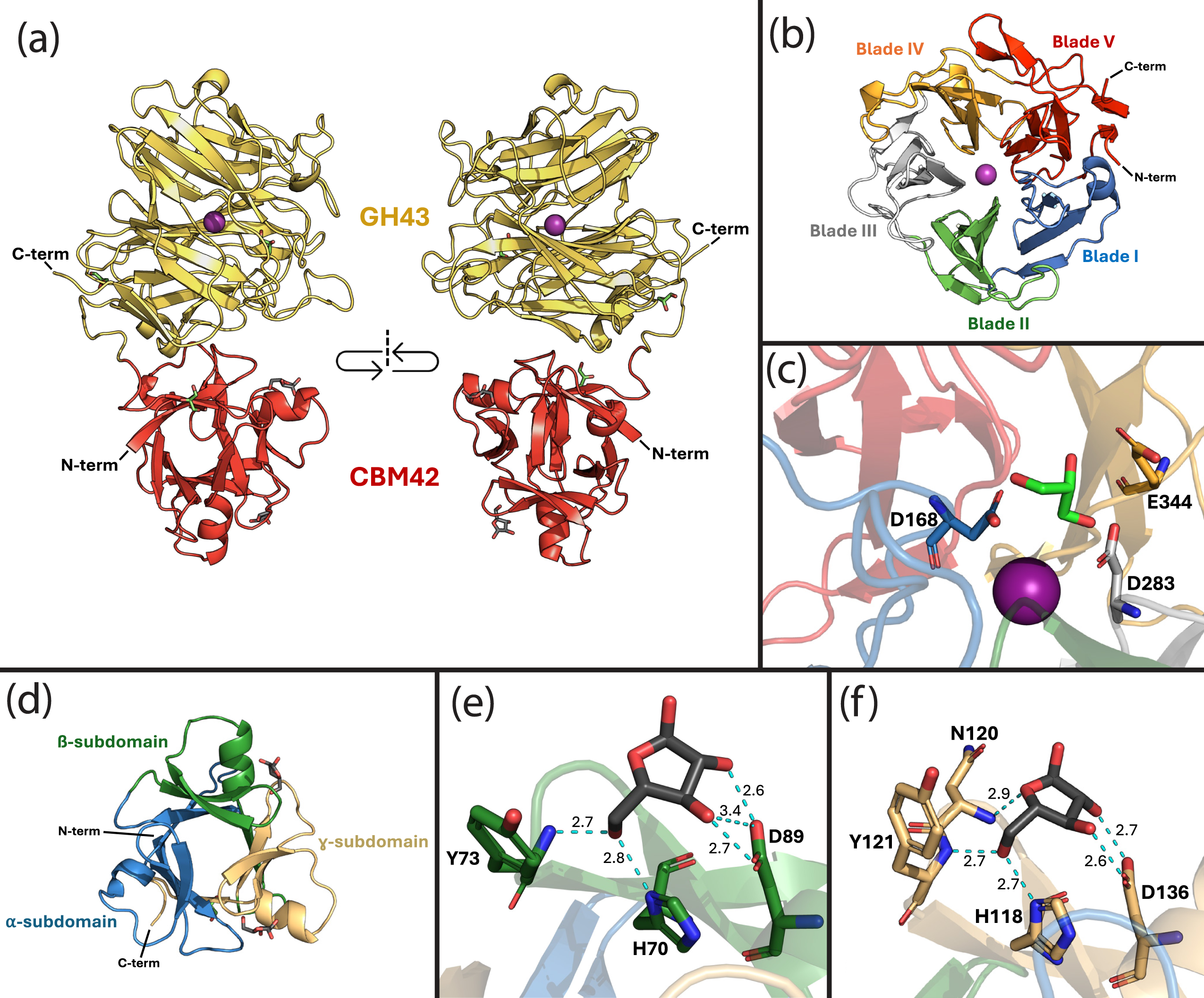
The major structural features of AtAbf43C shown using the arabinose-soaked structure (PDB code: 9NXH). Magnesium (Mg^2+^) is shown as a purple sphere, glycerol molecules are shown as green stick structures, and arabinose molecules are shown as dark grey stick structures. **(a)** The overall structure of AtAbf43C with the GH43 domain shown in yellow and the CBM42 domain shown in red. **(b)** The 5-bladed ß-propeller structure of the GH43 domain, with each blade shown in a distinct color. **(c)** The 3 catalytic residues labeled and shown as stick structures within the GH43 structure. **(d)** The ß-trefoil structure of the CBM42 domain, with each subdomain shown in a distinct color. **(e)** Arabinose bound in the ß-pocket of the CBM42 domain **(f)** Arabinose bound in the ɣ-pocket of the CBM42 domain. Contacted residues in **(e & f)** are labeled and shown as stick structures, Hydrogen bonds between the residues and arabinose molecule bonds are shown as dashed cyan lines, with bond lengths labeled in Å.

GH43 enzymes operate via an inverting mechanism in which three catalytic residues are highly conserved: aspartate acting as a base, a glutamate acting as an acid, and a second aspartate acting as a pKa modulator (9, 13, 24–30). In AtAbf43C these residues were identified to be D168, D283, and E344, found closely grouped at the base of the beta propeller structure (**Fig. 3c**.) To demonstrate the importance of these residues to catalytic activity, site-directed mutagenesis was used to generate versions of AtAbf43C (AtAbf43C_D168A, AtAbf43C_D283A, AtAbf43C_E344A, and AtAbf43C_H408A) in which each of the three active site residues, as well as histidine (H408) initially thought to interact with the magnesium ion, were individually mutated to alanine (Fig. S1a & b). When purified and tested alongside wild-type AtAbf43C, activity on pNPAra was completely eliminated in any of the versions of AtAbf43C where one of the three catalytic residues (D168, D283, and E344) was mutated to alanine (Fig. S1b; Table S7), indicating these sites are critical to enzymatic function. The H408A mutant appeared to have a small and statistically significant (p-value ≤ 0.05) increase in activity relative to the wild-type enzyme, suggesting this histidine residue is not critical for activity (Table S7).

The CBM42 domain of AtAbf43C displays the typical ß-trefoil structure found in other CBM42 and similar CBM13 family proteins consisting of three 40-50 amino acid subdomains (⍺, ß, and ɣ), each of which harbor a potential sugar binding pocket (**Fig. 3d**) (13, 14, 24). However, it has been observed in other CBM42 proteins that one of these three pockets may become nonfunctional (13, 14, 24). In the solved structure of arabinose-soaked Atabf43C_CBM42 (PDB 9NXH), arabinose was only found bound in the ß and ɣ pockets of the CBM42 domain (**Fig. 3d**). In the ß pocket the arabinose molecule formed hydrogen bonds with three residues, Y73, H70, and D89 (**Fig. 3e**). Similarly, the arabinose in the ɣ pocket contacted Y121, H118, and D136, as well as an additional residue N120 (**Fig. 3f**). In the previous study of this CBM42 by Ribeiro *et al*., residues D39, D91, and D136 were individually altered to alanine (14). While versions with the D91A and D136A substitutions showed significantly decreased binding affinity for arabinoxylan and arabinose relative to the wild-type protein, the D39A mutation, which corresponds to the ⍺ pocket, did not significantly affect binding (14). The binding pattern of arabinose observed in our structure would thus appear to support that the ⍺ pocket is non-functional or does not contribute to arabinose binding in the Atabf43C CBM42 domain.

Of the ten previously characterized GH43_26 enzymes (**Fig. 1b**; Table S2), SaAraf43A from the bacterium *Streptomyces avermitilis* is the only GH43_26 enzyme other than AtAbf43C with a crystal structure containing both the CBM42 and GH43 domains. Previously, SaAraf43A was extensively characterized by Ichinose *et al*. and Fujimoto *et al*., including multiple crystal structures (PDB: 3AKF, 3AKG, 3AKH, 3AKI) of the full-length protein complexed with various substrates, and a mutagenesis study identifying its catalytic residues (24, 31). SaAraf43A is an exo-1,5-⍺-L-arabinofuranosidase, with activity primarily towards AOSs such as arabinotriose, arabinotetraose, and arabinopentose (24, 31). Like AtAbf43C, SaAraf43A contains both a GH43 and CBM42 domain, but in SaAraf43C the GH43 domain resides at the N-terminus of the protein followed by a C-terminal CBM42 (24). Interestingly, despite the reverse ordering of these domains between SaAraf43A and AtAbf43C, the individual GH43 and CBM42 domains closely align. Structural alignment of the GH43 and CBM42 domains in AtAbf43C individually to the structure of arabinotriose complexed structure of SaAraf43A (PDB code: 3AKH) results in an RMSD of 0.475 Å (269 common C⍺ atoms) and 0.532 Å (111 common C⍺ atoms) respectively. (**Fig. 4a**; Fig S2a). As such, residues D168, D283, and E344 in the GH43 domain AtAbf43C closely aligned with corresponding catalytic residues D20, D135, and E196 in SaAraf43A (**Fig. 4b**), which when individually changed to alanine by Fujimoto *et al*. eliminated enzymatic activity on pNPAra (24). Furthermore, this structure of SaAraf43A contained an arabinose and arabinobiose molecule within the binding pocket of its GH43 domain. The topology of this binding pocket, characteristic of exo-acting GH43 enzymes, is such that the three catalytic residues are positioned at the bottom of a deep, narrow opening that sterically limits access to larger or branched substrates (24). This is opposed to endo-acting GH43s that are active on polysaccharides like arabinoxylan and arabinan, which possess a much more exposed binding cleft (24–28, 30). This narrowed binding pocket results from an extended loop structure in the 5^th^ blade of the beta propeller, which AtAbf43C appears to possess (**Fig. 3a** & b) (24, 25). When overlaid with the *apo* structure of the AtAbf43C GH43 domain, the bound arabinose and arabinobiose from SaAraf43A fit neatly within the surface structure of the AtAbf43C binding pocket (**Fig. 4c**), suggesting AtAbf43C maybe be active on similar AOS substrates as SaAraf43A. The binding domain of AtAbf43C also closely aligned with the CBM42 domain in SaAraf43A (Fig. S2a). However, while both structures had sugars in their ß-subdomains, the SaAraf43A CBM42 had an arabinobiose bound in its ⍺-pocket, and an unliganded ɣ-pocket (Fig. S2b-d). Furthermore, alignment of binding residues in the ⍺-pocket differed significantly between the two proteins, with AtAbf43C possessing a proline and two glutamines at positions where SaAraf43A possesses glutamine, histidine, and aspartate residues respectively (Fig. S2b). These differences could account for a non-functional ⍺-binding pocket in the AtAbf43C CBM42.

**Figure 4.**
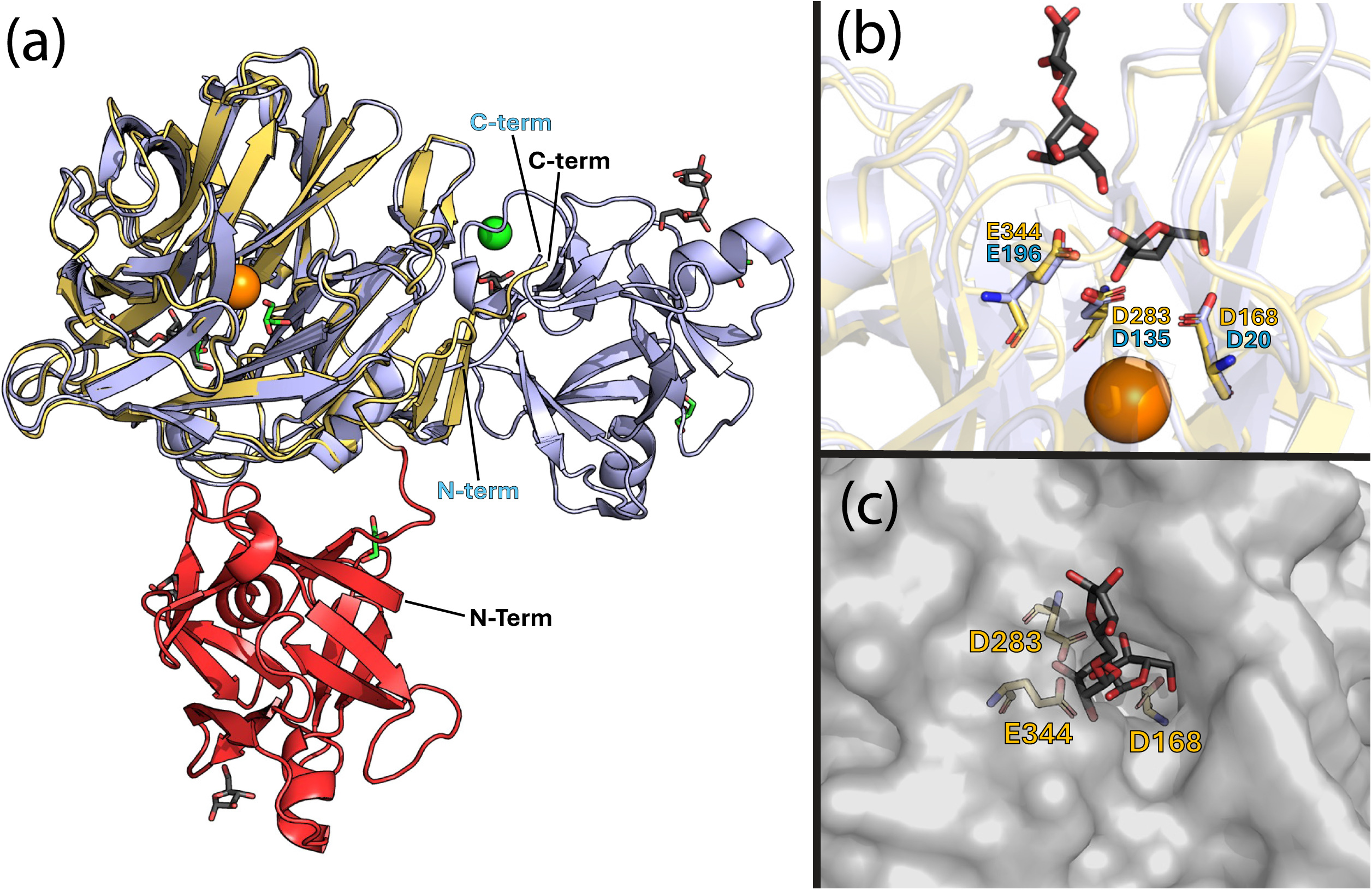
(a) Structural alignment of the GH43 domain of arabinose soaked AtAbf43C structure (GH43 domain in yellow, CBM42 domain in red) with the structure of SaAraf43A complexed with arabinotriose (PDB Code: 3AKH) shown in light blue. **(b)** Alignment of the catalytic residues in SaAraf43A (light blue) and co-crystalized arabinose and arabinobiose molecules in SaAraf43A structure with the active site residues of AtAbf43C (yellow). **(c)** Superimposition of the arabinose and arabinobiose molecules from the SaAraf43A structure onto the surface structure of AtAbf43C, with the AtAbf43C catalytic residues shown in yellow. In **(a-c)** sodium (Na^+^) is shown as an orange sphere, chlorine (Cl^-^) is shown as a green sphere, glycerol molecules are shown as green stick structures, and sugar molecules are shown as dark grey stick structures.

### LC-MS Analysis of Natural Polysaccharide and Oligosaccharide Hydrolysis

Finally, using liquid chromatography-mass spectrometry (LC-MS), the activity of AtAbf43C at optimal conditions was tested on the three natural substrates used previously (WAX, BX, SBA), as well as the following oligosaccharides: arabinobiose (A2), arabinotriose (A3), arabinotetraose (A4), arabinopentose (A5), 2^3^-α-L-arabinofuranosyl-xylotriose (A^2^XXX), and 3^3^-α-L-arabinofuranosyl-xylotetraose (XA^3^XXX). Based on these results, AtAbf43C appears to be primarily active towards α-1,5 linked AOS and to a lesser extent on arabinan where it seems to act in an exo manner to release free arabinose (**Table 3**; **Figure S3**). While marginal activity was detected on WAX, no significant activity was detected on BX or the AXOS tested, suggesting AtAbf43C does not degrade xylan-based substrates (**Table 3**).

**Table 3.** Summary of results from LC-MS analysis of AtAbf43C hydrolysis of various natural substrates and oligosaccharides. Mass spectra produced from the LC-MS based experiment and used for subsequent analysis can be found in the supporting information (**Fig. S3**).

| Substrate | WAX | BX | SBA | A2 | A3 | A4 | A5 | A<sup>2</sup>XXX | XA<sup>3</sup>XXX |
| --- | --- | --- | --- | --- | --- | --- | --- | --- | --- |
| Activity | +/- | - | + | + | + | + | + | - | - |

## Discussion

AtAbf43C is the fourth GH43 enzyme to be characterized from *A. thermocellus*, and only the third with an experimentally-determined structure (12, 17, 18, 29, 30). Furthermore, AtAbf43C is the first enzyme in the GH43_26 subfamily from *A. thermocellus*, and only the second crystal structure of a GH43_26 enzyme that includes the CBM domain (12, 17, 18, 24, 25) (**Table S2**). Natively AtAbf43C is likely an extracellular cellulosomal enzyme based on the presence of a N-terminal signal peptide and C-terminal dockerin I domain (4, 12, 17, 18). Characterization of the activity of AtAbf43C shows that it is an ⍺-arabinofuranosidase with activity towards pNPAra. The optimal pH and temperature for AtAbf43C on pNPAra (pH 5.5 and 65°C, **Fig 2a & b**) is generally consistent with other characterized *A. thermocellus* enzymes (4, 12, 17). Kinetic parameters determined on pNPAra show AtAbf43C had very high K_m_ as to approach the solubility limit of the substrate (**Table 1**), a result which was observed in the previously characterized *A. thermocellus* GH43 enzyme AxB8, an ⍺-arabinofuranosidase that was also primarily active on oligosaccharides (12). The larger *k_cat_*/*K_m_*value observed in the full length AtAbf43C protein versus the AtAbf43C_GH43 domain would suggest the CBM42 domain may aid in substrate specificity to pNPAra, however this is not definitive due to an inability to achieve substrate saturation due to the solubility limit of pNPAra. AtAbf43C was not active on the other pNP glycosides tested (**Table S5a**), indicating it acts primarily as an ⍺-arabinofuranosidase and does not have secondary function as a ß-xylosidase as has been reported for other GH43 enzymes (3–5, 12). Meanwhile, testing on natural hemicellulose substrates (WAX, BX, and SBA) using the DNS assay suggested that AtAbf43C was inactive towards xylan and arabinan polysaccharides (**Table S5b**).

Solution of multiple AtAbf43C crystal structures provided some insight into the enzyme’s preferred substrate and mode of action. First, the arabinose bound structure of full length AtAbf43C builds upon previous work by Riberio *et al*., by providing structural evidence for a non-functional ⍺-binding pocket, with arabinose only bound in the ß- and ɣ-subdomains (14). Subsequent alignment with the homologous CBM42 structure in SaAraf43A further supports this, as significant differences in corresponding binding residues were observed between the two proteins. Next, alignment of the GH43 domain in AtAbf43C with SaAraf43A allowed for identification of its three conserved active site residues in AtAbf43C (D168, D283, and E344), and insight into the binding modality of its binding pocket (24). Mutation of these catalytic residues in AtAbf43C confirms their involvement in its activity with all single point mutants losing activity (**Fig. S1, Table S7**). The deep-narrow topology of this pocket (**Fig 3c**), which limits access to the active site, is very similar to that of SaAraf43A (**Fig. 4b & c**), which acts in an exo manner towards ⍺-1,5-linked AOS. Subsequent LC-MS analysis of the hydrolysis products of AtAbf43C on natural substrates shows that this enzyme is capable of liberating some arabinose from SBA and, to a lesser extent, from WAX (**Table 3, Fig S3**), though notably not enough to be detected by the DNS assay, indicating that it likely acts in an exo manner at the ends of polysaccharides. LC-MS analysis of oligosaccharide hydrolysis (**Table 3, Fig S3**) showed AtAbf43C is active towards ⍺-1,5-linked AOS but not AXOS. This is consistent with activities observed in other members of the GH43_26 subfamily, which all act as arabinan- or AOS-degrading ⍺-1,5-arabinofuranosidases (**Table S2**). Taken together, we demonstrate through structural and functional characterization that AtAbf43C is an active arabinofuranosidase, specialized in degrading AOSs to arabinose in an exo manner. AtAbf43C from *A. thermocellus* is the most thermophilic GH43_26 enzyme characterized to date, thus deepening our understanding of this important subfamily of arabinofuranosidase enzyme.

## Experimental Procedures

### Sequence and Phylogenetic Analysis

The nucleotide and protein sequences for AtAbf43C (Locus tag: *Clo1313_2794*, GenBank Protein accession: ADU75776.1) as well as its predicted domains were found using the CAZy database and the National Center for Biotechnology Information (NCBI) database from the *A. thermocellus* DSM1313 genome (7, 32). The N-terminal signal peptide was predicted using the SignalP 6.0 software (33). The protein-protein BLAST search and pairwise alignment was conducted using the full amino acid sequence for Clo1313_2794 using the NCBI blastp tool. Characterized GH43 subfamily 26 proteins were found as annotated in the CAZy database, and their protein sequences were obtained using their primary ascension number in the NCBI database. Multiple sequence alignment of these proteins was performed using the ClustalOmega algorithm from which the phylogenetic tree was constructed using the IQTREE webserver and visualized using the Interactive Tree of Life (iTOL) online tool (34–36).

### Cloning of AtAbf43C

A table of oligonucleotide primers used to construct the plasmid in this study can be found in the supporting information (**Table S8)**. The gene for AtAbf43C was amplified from *A. thermocellus* DSM1313 genomic DNA purchased from the Leibniz Institute DSMZ-German Collection of Microorganisms and Cell Cultures via PCR (**Table S8;** Primers JLG005-006). Amplification removed a predicted N-terminal signal peptide and C-terminal dockerin I domain from the *Clo1313_2794* gene encoding the AtAbf43C module while adding overlap regions for subsequent Gibson assembly into the pET28b(+) expression vector (EMD Millipore) that added a C-terminal -LEHHHHHH purification tag. Truncated versions of AtAbf43C (AtAbf43C_CBM42 and AtAbf43C_GH43) were amplified from this resulting vector with overlaps for Gibson assembly into the pET28b(+) vector (**Table S8**; Primers JLG005 & JLG036-38). Site-directed mutagenesis of AtAbf43C was carried out via PCR amplification of the full-length expression plasmid using mutagenic primers which added overlap regions for subsequent recirculation via Gibson assembly (**Table S8**; Primers JLG195-202). Gibson Assemblies were performed by incubating amplified DNA fragments at 50 ℃ for 1 hour with NEBuilder HiFi DNA Assembly Master mix (New England Biosciences) as per the manufacturer’s recommended protocol.

### Bacterial Strains and Culture Conditions

Plasmids were cloned in NEB chemically competent *Escherichia coli* DH5⍺ (New England Biolabs). Plasmids were isolated using ZymoPURE miniprep kits (Zymo Research), and plasmid sequences were confirmed by sequencing (Azenta Genewiz). Sequenced confirmed plasmids were then transformed into chemically competent *E. coli* BL21 (DE3) pRosetta2 (EMD Millipore) for protein expression. *E. coli* cultures were maintained in enriched Luria-Bertani (LB) Medium (24 g/L yeast extract, 10 g/L tryptone, 5 g/L NaCl) or LB medium (5 g/L yeast extract, 10 g/L tryptone, 5 g/L NaCl, 15 g/L agar) (1.5% w/v) agar plates with 50 µg/ml kanamycin (IBI Scientific), or 50 µg/ml kanamycin (IBI Scientific) and 33 µg/ml chloramphenicol (RPI), as appropriate.

### Protein Expression and Purification

Protein expression was induced by inoculating ZYM-5052 autoinduction media with overnight cultures of the transformed *E. coli* BL21 DE3 Rosetta strains (37). Cells were harvested by centrifugation at 6000 x g for 10 minutes after 18-22 hours of growth at 37 ℃ in a shaking incubator at 250 rpm. Cell pellets were resuspended in lysis buffer (20 mM Sodium Phosphate pH 7.4, 500 mM NaCl, 10 mM imidazole) before being lysed via sonication on ice using a Branson SFX 550 Sonifer® in cycles of 10s on at 20 kHz and 10s off for 10 minutes total. This lysate was then centrifuged for 30 minutes at 30,000 x g at 4 ℃, and the resulting supernatant was passed through a 0.22 µm filter. Full length AtAbf43C and its truncated versions were then purified via Immobilized Metal Affinity Chromatography (IMAC) using 5 ml EconoFit IMAC columns (Bio-Rad) on an NGC Chromatography System (Bio-Rad) and fractionated in elution buffer (20 mM Sodium Phosphate pH 7.4, 500 mM NaCl, 250 mM imidazole). Fractions of IMAC purified full length AtAbf43C protein were combined and further purified via size exclusion chromatography via FPLC using a HiLoad^TM^ 26/600 Superdex^TM^ 200 pg. column (Cytiva) in a mobile phase of 50 mM sodium phosphate pH 7.0, 150 mM NaCl buffer. Mutant versions of AtAbf43C and wild type AtAbf43C protein used to test the effect of mutagenesis were purified via immobilized metal ion chromatography using His-Spin Protein Mini Prep Kits (Zymo Research) as per the manufacturer’s instructions. After purifications, all proteins were buffer exchanged into pH 6.0 100 mM sodium phosphate buffer using 30 kDa MWCO Centrifugal Filter Units (CELLTREAT®) or 10 kDa MWCO Spin-X® UF 20 mL Centrifugal Concentrators (Corning®). Purity of the resulting proteins was assessed by SDS-PAGE using 4–20% Mini-PROTEAN® TGX Stain-Free^TM^ Protein Gels (Bio-Rad) with Precision Plus^TM^ unstained protein standards (Bio-Rad). Protein concentration was evaluated by measuring the A280nm of the resulting buffer exchanged proteins using a Nanodrop One spectrophotometer (Thermo Scientific) and calculating protein concentration using Beer’s law and the calculated A280nm extinction coefficient of each individual protein (38–40). Aliquots of full length AtAbf43C were frozen at -80°C prior to further analysis at a protein concentration of 9mg/mL.

### Substrates

Para-nitrophenol-⍺-L-arabinofuranoside (pNPAra) and para-nitrophenol-ß-D-xylopyranoside (pNPXy) were obtained from Sigma-Aldrich and EMD Millipore respectively. para-nitrophenol-⍺-D-galactopyranoside (pNP⍺Gal) and para-nitrophenol-ß-D-galactopyranoside (pNPßGal) were obtained from TCI chemicals. Natural substrates with purities >95% including Wheat Arabinoxylan (WAX), Beechwood Xylan (BX), and Sugar Beet Arabinan (SBA) were obtained from Megazyme (Neogen). Oligosaccharides including Arabinobiose, Arabinotriose, Arabinotetraose, Arabinopentose, 2^3^-α-L-Arabinofuranosyl-xylotriose (A^2^XXX), and 3^3^-α-L-Arabinofuranosyl-xylotetraose (XA^3^XXX) were obtained from Megazyme (Neogen). Arabinose and Xylose were purchased from Fisher Scientific.

### Temperature and pH optimization

Prior to all assays AtAbf43C protein at a final concentration of 9 mg/ml in pH 6 100 mM sodium phosphate buffer was diluted to 0.025 mg/ml in an appropriate reaction buffer, described further below. Reactions were initiated by adding 45 µl of 5 mM pNPAra solution dissolved in appropriate buffer to 5 µl of the 0.025 mg/ml enzyme solution in PCR strip tubes, before immediately being moved to a thermocycler for incubation. After 10 minutes, reactions were stopped with the addition of 100 µl of 1M sodium carbonate. The absorbance at 405 nm of 100 µl of each reaction were measured in a flat-bottomed clear 96 well plate using a BioTek SynergyH1 microplate reader (Agilent). pH optimization was first performed at 55 ℃ with 100 mM sodium acetate buffer used for pH 4-5.5 conditions, and 100 mM sodium phosphate buffer for pH 6-8 conditions. Temperature optimization was then performed at the optimal observed pH of 5.5 in 100 mM sodium acetate buffer. Activity was calculated as a percentage relative to the highest observed activity. All reaction conditions were performed in triplicate.

### Thermostability and Effect of Additives

To test the thermostability of AtAbf43C, protein was first diluted as described previously in pH 5.5 100 mM sodium acetate buffer, before being incubated in a thermocycler at temperatures of 55-70 ℃. Aliquots of protein were then removed at 30-minute, 1-hour, 3-hour, and 6-hour timepoints. Activity was then tested as described above at the optimal observed temperature of 65 ℃ using 5 mM pNPAra dissolved in pH 5.5 100mM sodium acetate buffer. Residual activity was calculated as a percentage relative to that of unincubated AtAbf43C. To test the effects of various salts and chelating agents, AtAbf43C was first diluted in pH 5.5 100 mM sodium acetate buffer containing the additive at 10 mM and pre-incubated at room temperature for 1-hour prior to adding substrate. Activity was then tested as described previously at optimal temperature and pH except the 5 mM pNPAra substrate solutions also contained 10 mM of the specific additive. Additives were as follows: NaCl, CaCl_2_, KCl, MgSO_4_*7H_2_O, MnCl_2_*4H_2_O, ZnSO_4_*7 H_2_O, CuCl2*2H_2_O, FeCl_3_*6H_2_O, CoCl_2_, NiCl_2_*6H_2_O, EDTA tetrasodium dihydrate. Activity was calculated as a percentage relative to AtAbf43C incubated and tested with no additive. Additionally, the absorbance at 405nm measured from blank solutions containing additives were subtracted from those observed in the corresponding enzymatic reaction conditions to control for the variation in absorbance due to the addition of the specific metal salt or chelating agent. All reaction conditions were performed in triplicate. Statistical significance of additive effects was determined by running a Brown-Forsythe and Welch ANOVA test on the collected data in GraphPad Prism version 10.0 (GraphPad Software, Boston, Massachusetts USA).

### Substrate Specificity

Activity against pNP-Glycosides (pNPAra, pNPXy, pNP⍺Gal, and pNPßGal) was tested at optimal pH and temperature as described above for pNPAra on AtAbf43C, AtAbf43C_GH43, and AtAbf43C_CBM42. 4-nitrophenol (pNP) released was quantified via Beer’s law with a 405 nm extinction coefficient of 18500 L/(mol*cm) and path length calculated empirically as per the manufacturer’s recommendation (41, 42). Specific activity was then calculated as U/mg of protein, where U is defined as µM/s of released pNP. Activity against natural substrates Wheat Arabinoxylan (WAX), Beechwood Xylan (BX), and Sugar Beet Arabinan (SBA), was then tested by incubating the proteins as described above at optimal conditions except with substrate solutions containing natural substrate dissolved at 1% (w/v) in buffer and incubation time lengthened to 1 hour. Activity was detected as described previously by Conway *et al.* using the dinitrosalicylic acid (DNS) reducing assay with L-arabinose used as a standard for oligosaccharide release (43, 44).

Kinetic parameters were determined for AtAbf43C and AtAbf43C_GH43 by incubating the proteins as described above except with varying concentrations of pNPAra (0.9-36 mM) and with reaction times shorted to 2 minutes to ensure linear initial rates of reaction. pNP released was quantified as described above with velocities calculated as µM pNP/s. Using the predicted molar mass of each protein, kinetic parameters were then calculated using the predicted molar mass via non-linear regression using the “determine kcat” model in GraphPad Prism version 10.0 (GraphPad Software, Boston, Massachusetts USA).

### Testing of AtAbf43C Mutants

To test the importance of certain residues to catalytic activity, purified AtAbf43C with residues D168, D283, E344, and H408 individually mutated to alanine were tested alongside wild type AtAbf43C on pNPAra. Proteins were prepared and incubated in pH 5.5 sodium acetate with 5 mM pNPAra substrate as described above for 10 minutes at 65 ℃ at a final reaction concentration of 0.0025 mg/ml protein.

Activity was calculated as a percentage relative to that of wild type AtAbf43C. Statistical significance was determined by running a Brown-Forsythe and Welch ANOVA test on the collected data in GraphPad Prism version 10.0 (GraphPad Software, Boston, Massachusetts USA).

### LC-MS Analysis of Hydrolyzed Products

To further investigate the activty of AtAbf43C, reactions were initiated by adding 90 µL of substrate solution dissolved in pH 5.5 100 mM sodium acetate buffer to 10 µL of enzyme diluted in the same buffer as described above at 0.025 mg/ml or 10 µL of blank buffer. Samples were incubated at 65 ℃ for 1 hour in a thermocycler after which the reactions were stopped by heating at 95 ℃ for 5 minutes to inactivate the enzyme. Natural substrates WAX, BX, and SBA were dissolved at a concentration of 1% (w/v), while oligosaccharides and sugars were dissolved at 0.1% w/v. LC-MS analysis was performed using an Agilent 6530 QTOF connected to an Agilent 1260 LC system. Mass spectra were acquired using electrospray ionization (ESI) with the instrument in positive ion mode. Reaction samples were run on a Agilent HI-PLEX Na (Octo) 300 x 7.7 mm column heated to 80 °C. The mobile phase was ultrapure water, and sample runs were 30 minutes long with a flow rate of 0.5 mL/min. Data were analyzed using Agilent MassHunter software; mass spectra and extracted ion chromatograms (EICs) for species of interest ([M+Na]^+^ adducts) were then obtained.

### Crystallization, data collection, and structure refinement

Purified AtAbf43C, AtAbf43C_CBM42, and AtAbf43C_GH43 proteins at concentrations of 9.0 mg/ml, 4.4 mg/ml, and 8.5 mg/ml respectively were crystalized over the course of several days via the sitting drop vapor diffusion method. AtAbf43C was crystalized in the form of thin needle-like blades in space group P2_1_ with two molecules in the asymmetric unit from a solution of 15-20% v/v propanol, 25% w/v PEG3350, 50 mM AmSO_4_, and 0.1 M HEPES pH 7.7. AtAbf43C crystals soaked in L-arabinose, crystallized in similar conditions, were in space group C2 with one molecule in the asymmetric unit. AtAbf43C_CBM42 was crystalized in space group P2_1_2_1_2 with one molecule in the asymmetric unit in the form of small flat plates from a solution of 25% w/v PEG3350 and 0.1 M citric acid pH 3.0. AtAbf43C_GH43 was crystalized in space group P6_5_ with two molecules in the asymmetric unit in the form of bi-pyramidal crystals in 25% w/v polyethylene glycol monomethyl ether (pegM) 5000, 0.25 M AmS04, and 0.1 M Bis-Tris pH 5.5. Crystals were mounted in nylon loops (Hampton Research) and flash-cooled in liquid nitrogen after brief (<30 seconds) equilibration in cryoprotectant solutions. Cryoprotectant solutions corresponded to crystallization solutions supplemented with 27-30% v/v glycerol. For the arabinose soak of AtAbf43C crystals the cryoprotectant solution was also supplemented with 5% w/v arabinose and the crystal equilibration time in this solution was extended to 3 minutes before flash-cooling.

Data were collected on beam lines 17-ID-1 (AMX) and 17-ID-2 (FMX) at Brookhaven National Lab, Upton, New York, USA. Data were processed using XDS and scaled using AIMLESS (45, 46). Data collection and processing statistics are shown in **Table 2**. Structures of AtAbf43C, AtAbf43C_CBM42, and AtAbf43C_GH43 proteins were determined by molecular replacement using the program PHASER starting from the AlphaFold model of AtAbf43C and partially refined versions thereof (47). AtAbf43C_GH43 was first solved using an edited AlphaFold model, and the partially refined structure was used as a model for AtAbf43C which was then completed using the AlphaFold model for AtAbf43C_ CBM42. Partial refinement of this full-length AtAbf43C model at 1.3 Å resolution was then used as the starting point for the solution of AtAbf43C_CBM42. Initial models were iteratively rebuilt and refined using COOT and PHENIX.REFINE respectively (47, 48). An apparent metal binding site was added based on local site geometry and surrounding ligands and assigned as Mg^2+^. Arabinose binding to AtAbf43C was assessed by model-phased difference (||Fobs|-|Fcalc||) maps revealing two arabinose molecules in the furanose conformation bound to AtAbf43C. We were not successful in obtaining arabinose bound to AtAbf43C_GH43 in soaking experiments. Persistent difference density at the expected active site of AtAbf43C_GH43, too large to be the cryoprotectant used, was modeled using a carbohydrate group formally named as an unknown ligand (residue name UNL). No oligomerization of the AtAbf43C, AtAbf43C_CBM42, and AtAbf43C_GH43 proteins were observed within the crystal lattices of the four crystal forms. Final refinement statistics are shown in **Table 2**. The four structures have been deposited in the Protein Data Bank (AtAbf43C with id 9NXG, AtAbf43C:Arabinose with id 9NXH, AtAbf43C_CBM42 with id 9NXI, and AtAbf43C_GH43 with id 9NXJ (49). Visualizations were prepared and structural alignments were performed using the PyMOL Molecular Graphics System, Version 2.5.8 Schrödinger, LLC.

## Data Availability

Protein crystal structures obtained in our study are deposited to the Protein Data Bank with accession numbers 9NXG, 9NXH, 9NXI, 9NXJ. All other data is contained within the manuscript and supplementary information.

## Supporting Information

This article contains supporting information.

## Acknowledgements

This research used resources of the National Synchrotron Light Source II, a U.S. Department of Energy (DOE) Office of Science User Facility operated for the DOE Office of Science by Brookhaven National Laboratory under Contract No. DE-SC0012704. The Center for BioMolecular Structure (CBMS) is primarily supported by the National Institutes of Health, National Institute of General Medical Sciences (NIGMS) through a Center Core P30 Grant (P30GM133893), and by the DOE Office of Biological and Environmental Research (KP1605010).

## Funding and additional information

This work was supported by the Energy Research Fund administered by the Andlinger Center for Energy and the Environment at Princeton University and startup funds from the Department of Chemical and Biological Engineering at Princeton University to J.M.C. Mass spectrometry data were collected on an instrument purchased with a supplement to NIH grant GM107036 to A.J.L. A.Z. was supported by an NSF Graduate Research Fellowship Program under Grant DGE-2039656.

## Abbreviations and nomenclature

(CAZymes): Carbohydrate Active enZymes
(GH): Glycoside hydrolase
(CBM): Carbohydrate Binding Module
(AOS): arabino-oligosaccharides
(AXOS): arabino-xylo-oligosaccharides
(pNP): 4-nitrophenol
(pNPAra): para-nitrophenol-α-L-arabinofuranoside
(pNPXy): para-nitrophenol-ß-D-xylopyranoside
(pNP⍺Gal): para-nitrophenol-⍺-D-galactopyranoside
(pNPßGal): para-nitrophenol-ß-D-galactopyranoside
(DNS): dinitrosalicylic acid
(WAX): wheat arabinoxylan
(BX): beechwood xylan
(SBA): sugar beet arabinan
(A2): Arabinobiose
(A3): Arabinotriose
(A4): Arabinotetraose
(A5): Arabinopentose
(A^2^XXX): 2^3^-α-L-Arabinofuranosyl-xylotriose
(XA^3^XXX): 3^3^-α-L-Arabinofuranosyl-xylotetraose

