## Supplementary Figures for "Functional and structural characterization of AtAbf43C: An exo-1,5-⍺-L-arabinofuranosidase from *Acetivibrio thermocellus* DSM1313"

**Figure S1.** **(a)** Structural visualization of residues mutated in AtAbf43C in the arabinose-soaked structure (PDB: 9NXH). **(b)** Protein gel of expressed and purified wild-type and mutant AtAbf43C used in subsequent activity assays.

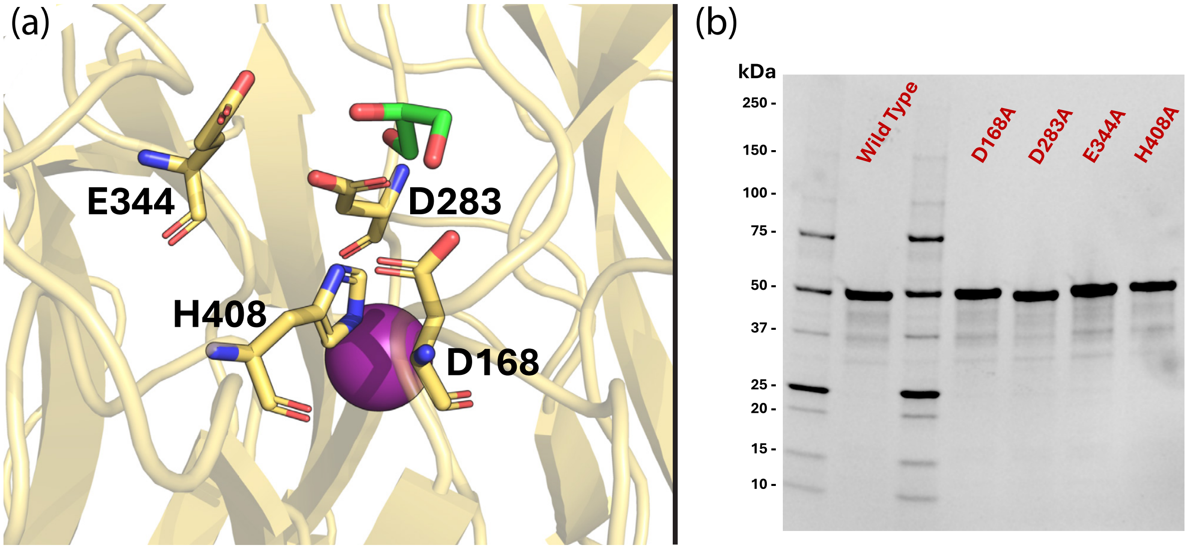

**Figure S2. (a)** Structural alignment of the CBM42 domain of arabinose soaked AtAbf43C structure (GH43 domain in yellow, CBM42 domain in red) with the structure of SaAraf43A complexed with arabinotriose (PDB Code: 3AKH) shown in light blue. **(b)** Alignment of residues that form hydrogen bonds with the arabinobiose molecule bound in α-pocket of the SaAraf43A CBM42 domain (light blue) with corresponding residues in the Atabf43C structure (red). **(c)** Alignment of residues that form hydrogen bonds with the arabinobiose molecule bound in ß-pocket of the SaAraf43A CBM42 domain (light blue) with corresponding arabinose bound residues in the Atabf43C structure (red). **(d)** Alignment of residues that form hydrogen bonds with the arabinose molecule bound in ɣ-pocket of the AtAbf43C CBM42 domain (red) with corresponding residues in the SaAraf43A structure (light blue). In **(a-d)** magnesium ($Mg^{2+}$) is shown as an orange sphere, sodium ($Na^{+}$) is shown as an orange sphere, chlorine ($Cl^{-}$) is shown as a green sphere, glycerol molecules are shown as green stick structures, and sugar molecules are shown as dark grey stick structures. Hydrogen bonds between residues and sugars the SaAraf43A structure are shown as dashed cyan lines, while hydrogen bonding in the AtAbf43C structure are shown as yellow dashes.

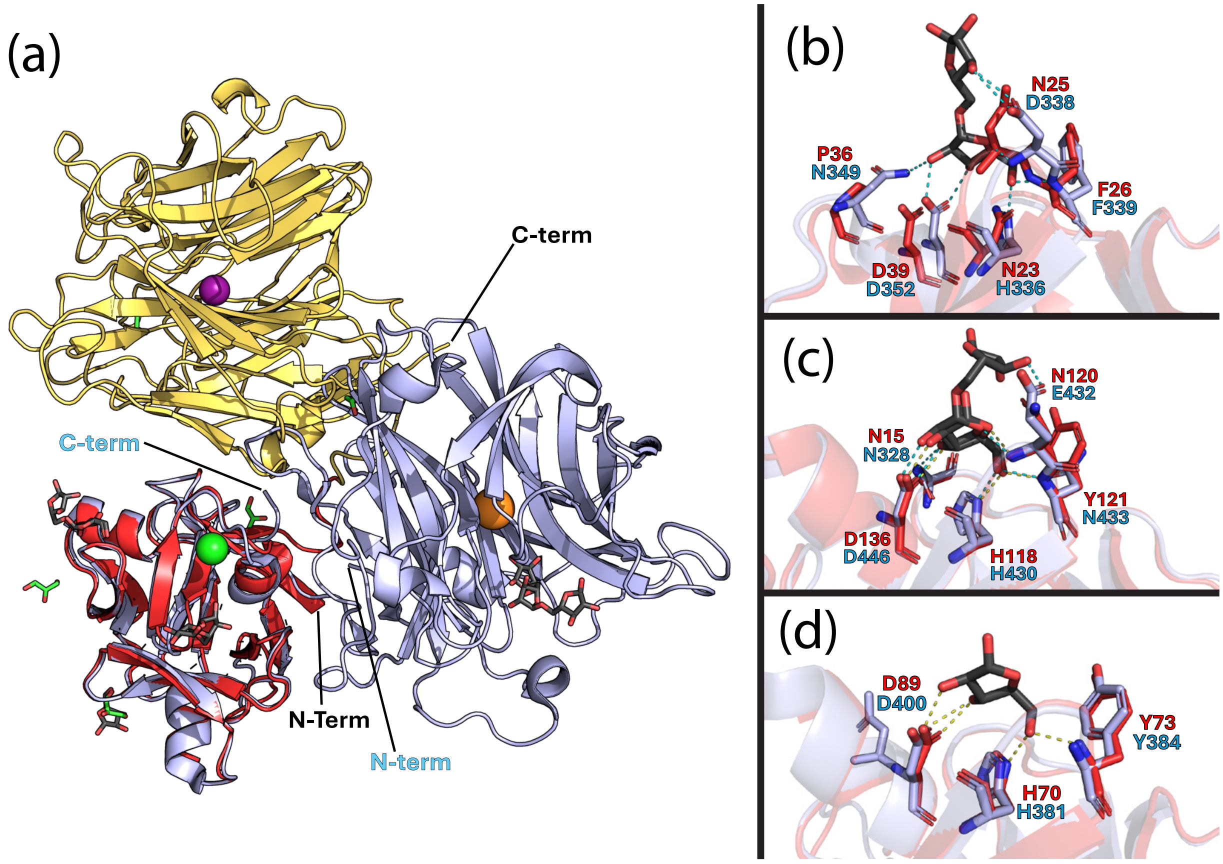

**Fig S3.** Mass spectra for AtAbf43C *in vitro* hydrolysis reactions on natural polysaccharides and oligosaccharides. **(a)** Table of retention times and visual key for relevant species. Reactions were performed without and with AtAbf43C on **(b)** arabinose, **(c)** xylose, **(d)** WAX, **(e)** BX, **(f)** SBA, **(g)** A2, **(h)** A3, **(i)** A4, **(j)** A5, **(k)** A2XXX, and **(l)** XA3XXX. For each reaction, a mass spectrum was acquired for a range of 8 to 18 min in retention time, as all products of interest have retention times within this range. All species were identified as $[M+Na]^+$ adducts.

**(a)**

| Molecule | Formula | MW | (M+Na)+ | Retention time of center of peak |
|---|---|---|---|---|
| Arabinose | C5H10O5 | 150.13 | 173.042 | ~15.85 min |
| Xylose | C5H10O5 | 150.13 | 173.042 | ~14.6 min |
| A2 | C10H18O9 | 282.2 | 305.0843 | ~11.6 min |
| A3 | C15H26O13 | 414.4 | 437.1266 | ~10.35 min |
| A4 | C20H34O17 | 546.5 | 569.1688 | ~9.7 min |
| A5 | C25H42O21 | 678.6 | 701.2111 | ~9.35 min |
| A-2-XXX | C20H34O17 | 546.5 | 569.1688 | ~9.9 min |
| X-A-3-XXX | C25H42O21 | 678.6 | 701.2111 | ~9.6 min |

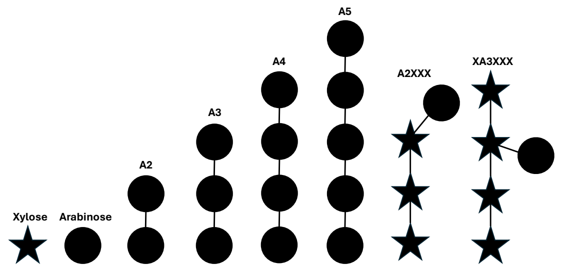


**(b)**

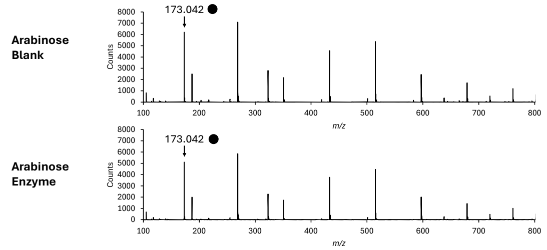


**(c)**

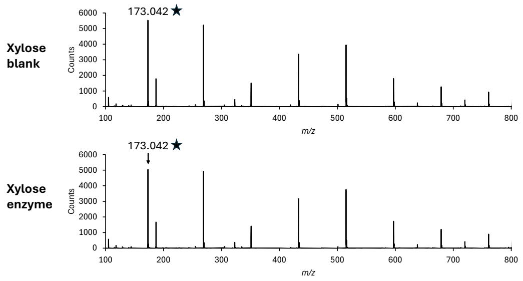


**(d)**

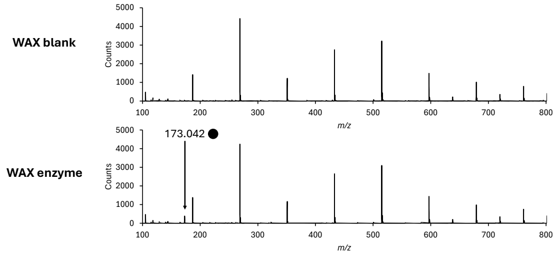


**(e)**

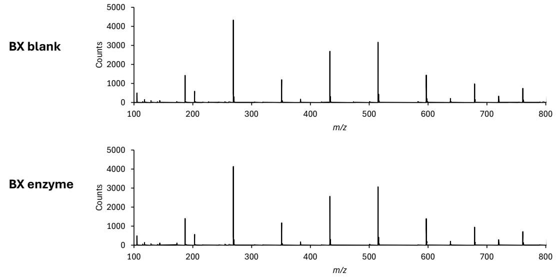


**(f)**

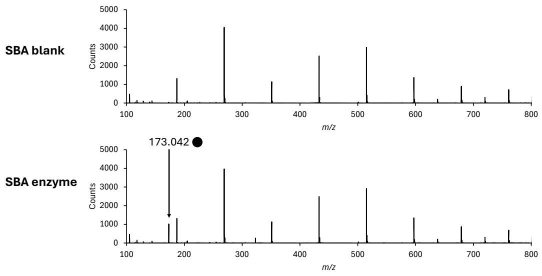

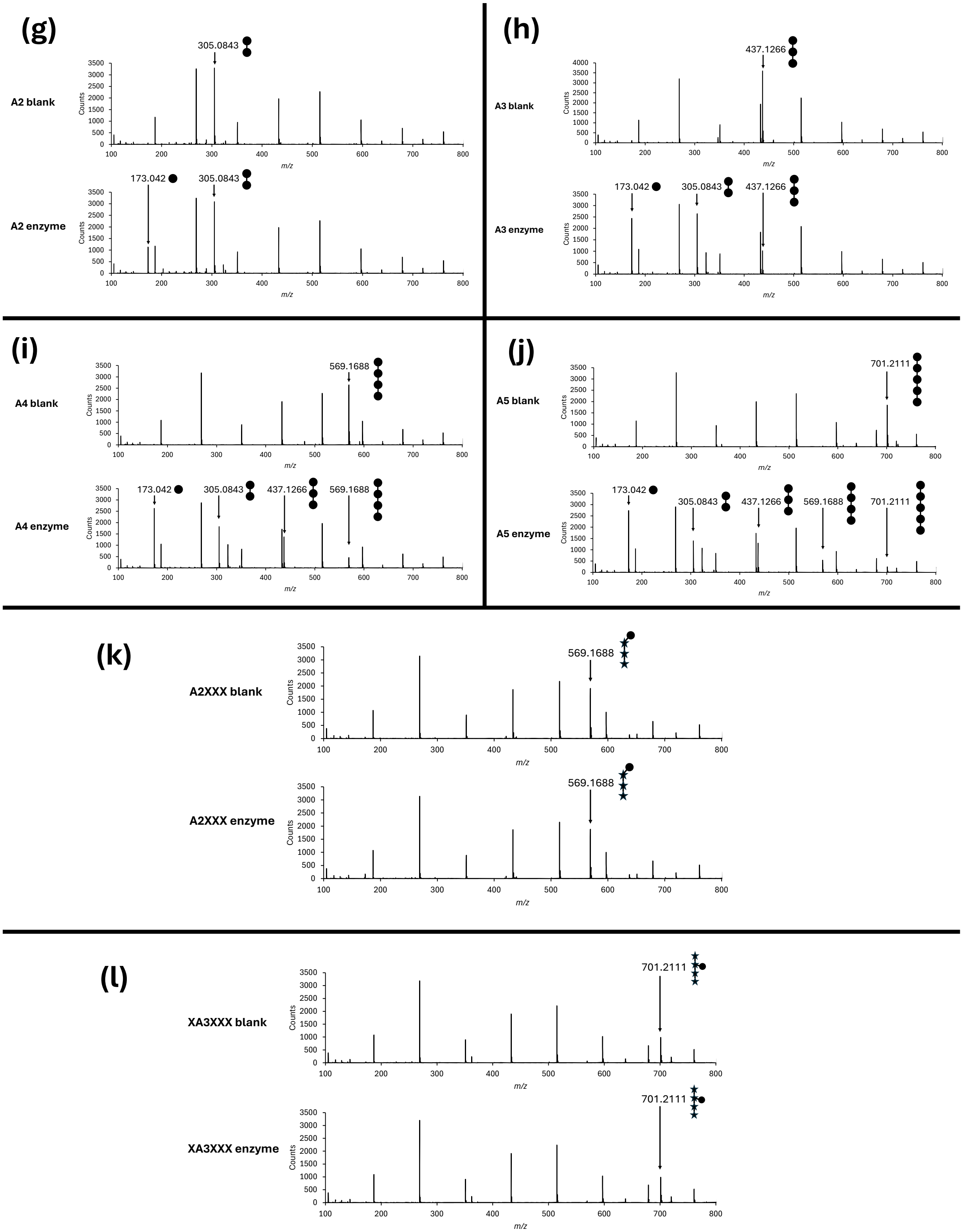
(g)
A2 blank
305.0843
A2 enzyme
173.042
305.0843
(h)
A3 blank
437.1266
A3 enzyme
173.042
305.0843
437.1266
(i)
A4 blank
569.1688
A4 enzyme
173.042
305.0843
437.1266
569.1688
(j)
A5 blank
701.2111
A5 enzyme
173.042
305.0843
437.1266
569.1688
701.2111
(k)
A2XXX blank
569.1688
A2XXX enzyme
569.1688
(l)
XA3XXX blank
701.2111
XA3XXX enzyme
701.2111
Counts
m/z
